# Tissue-Specific and Interorgan Metabolic Reprogramming Maintains Tolerance to Sepsis

**DOI:** 10.1101/2022.10.08.511411

**Authors:** Brooks P. Leitner, Won D. Lee, Wanling Zhu, Xinyi Zhang, Rafael C. Gaspar, Zongyu Li, Joshua D. Rabinowitz, Rachel J. Perry

## Abstract

Reprogramming metabolism is of great therapeutic interest for reducing morbidity and mortality during sepsis-induced critical illness^1^. Disappointing results from randomized controlled trials targeting glutamine and antioxidant metabolism in patients with sepsis have begged for both identification of new metabolic targets, and a deeper understanding of the metabolic fate of glutamine at the systemic and tissue-specific manner^2–4^. In critically ill patients versus elective surgical controls, skeletal muscle transcriptional metabolic reprogramming is comprised of reduced expression of genes involved in mitochondrial metabolism, electron transport, and glutamate transport, with concomitant increases in glutathione cycling, glutamine, branched chain, and aromatic amino acid transport. To analyze putative interorgan communications during sepsis, we performed systemic and tissue specific metabolic phenotyping in a murine polymicrobial sepsis model, cecal ligation and puncture. In the setting of drastically elevated inflammatory cytokines, we observed >10% body weight loss, >50% reductions in oxygen consumption and carbon dioxide production, and near full suppression of voluntary activity for the 48 hours following sepsis as compared to sham-operated controls. We found increased correlations in the metabolome between liver, kidney, and spleen, with drastic loss of correlations between the heart and quadriceps metabolome and all other organs, pointing to a shared metabolic signature within vital abdominal organs, and unique metabolic signatures for skeletal and cardiac muscle during sepsis. A lowered GSH:GSSG and elevated AMP:ATP ratio in the liver underlie the significant upregulation of isotopically labeled glutamine’s contribution to TCA anaplerosis and glutamine-derived glutathione biosynthesis; meanwhile, the skeletal muscle and spleen were the only organs where glutamine’s contribution to the TCA cycle was significantly suppressed. These results highlight tissue-specific mitochondrial reprogramming, rather than global mitochondrial dysfunction, as a mechanistic consequence of sepsis. Using a multi-omic approach, we demonstrate a model by which sepsis-induced proteolysis fuels the liver’s production of anaplerotic substrates and the antioxidant glutathione to sustain tolerance to sepsis.

## Introduction

Sepsis-induced critical illness affects millions of individuals in the US and worldwide every year^5^. Even among survivors, a large fraction experience post-septic complications, with a higher-than-expected proportion of sepsis survivors either dying or returning to hospitals within the next year compared to an age-matched population^6,7^. The acuity of life-threatening sepsis and the existence of highly reproducible metabolic shifts during both the hyper- and hypometabolic phases suggests that metabolic modulation may be a promising avenue for therapy. In the past decade, however, numerous trials attempting to supplement critically ill patients with various metabolic modulators, including selenium, glutamine, vitamin C, and thiamine^3,8–10^, have yielded minimal survival benefit if not detrimental as in the glutamine supplementation^2^. This has led critical care leaders of the International Surviving Sepsis Campaign to prioritize a better understanding of the cellular and subcellular mechanisms of sepsis-induced metabolic reprogramming^1^.

Sepsis is a condition of disrupted inflammation-initiated metabolic homeostasis. Corresponding to inflammatory response changes, sepsis develops from early (hypermetabolic) to late (hypometabolic) phase. During the early hypermetabolic phase, muscle proteolysis and liver gluconeogenesis increase to provide fuel for fighting against infection and healing wounds. Afterward, during the hypometabolic phase, the whole-body shifts to fatty acid and amino acid oxidation to resist organ damage. Finally, a longer-duration phase follows that corresponds to tolerance to sepsis^11–13^. The hypermetabolic phase is notoriously challenging to identify in patients given the heterogeneity of presentation, differences in sepsis onset prior to or while in the intensive care unit or emergency room, in addition to variance in currently poorly defined patient characteristics^14^. Further, the characterization of damage to muscle and vital organs during the hypometabolic phase is also poorly understood, with several mechanisms describing the liver’s role in coordinating metabolic reprogramming of lipid metabolism and iron metabolism during this time^15–17^. Consequently, there is an urgent need for further understanding of the factors related to systemic metabolic reprogramming during sepsis.

Given that critical illness-induced skeletal muscle wasting has been thoroughly documented, and likely contributes both to mortality and long-term morbidity, we began our investigation with transcriptomic analyses of human skeletal muscle biopsies taken in the intensive care unit. Then, given that evolutionary pressures have required limited wasting of precious amino acids, we reasoned that the constituent amino acids derived from skeletal muscle likely are destined to contribute to other more life-threatening functions. We then performed targeted and untargeted plasma and tissue metabolomics after metabolic characterization of our cecal ligation and puncture (CLP) murine model of polymicrobial sepsis with a comprehensive cytokine panel and metabolic cage studies. Finally, using insight from our discovery-based transcriptomic and metabolomic analyses, we performed stable isotope tracer infusion studies with labelled glutamine and glucose to examine the metabolic mechanisms of sepsis-induced metabolic reprogramming in seven tissues and plasma during sepsis.

Using integrated systems biology approaches, we uncover shared metabolic patterns in the resting state of the organism, and the tissue-specific relationships in response to sepsis. In the resting state, S-adenosylmethionine, which largely serves as a broad housekeeping metabolite for methyl transfers during amino acid synthesis and interconversion, nucleotide regulation, and other pathways, is the most significantly correlated metabolite across all organs. During sepsis, nearly all inter-tissue connections are lost, while glutamine and glutamate become the most significantly correlated metabolites in all tissues except for cardiac and skeletal muscle. We thus reveal shared and tissue specific metabolic responses in the resting state and in response to sepsis, a unique demonstration of systemic metabolic plasticity.

We believe that the integrative approach taken here provides a metabolic mechanism for the destiny of glutamine and glutathione at the systemic level during sepsis, and identifies new therapeutic leads for modulating metabolism during sepsis. A better characterization of the shared and tissue-specific metabolic response to sepsis provides greater resolution for metabolic-targeted therapy.

## Results

### Septic Patient Skeletal Muscle Transcriptomics Reveals Proteolysis, Electron Transport Chain Dysfunction, and Upregulation of Amino Acid Transport

Skeletal muscle biopsies (vastus lateralis) were taken during a previous study in 13 critically ill septic patients currently in the intensive care unit (ICU) and 8 healthy control patients who were undergoing elective surgical procedures with confirmed absence of infection and the samples were subjected to transcriptomic analysis^18^. Thousands of genes were differentially expressed, and notably the gene encoding myosin heavy chain 7B, a structural component of type I oxidative muscle fibers, was downregulated greater than 32 fold compared to control skeletal muscle, suggesting that sepsis induces a major defect in the oxidative muscle (Fig 1A). Significantly upregulated genes included the glutamine transporter SLC38A1 and glutaredoxin, suggesting reprogramming of glutamine and redox metabolism within skeletal muscle. To characterize metabolites the muscle may prioritize importing or exporting during sepsis, we isolated all solute carrier (SLC) genes (Fig 1B). The most upregulated SLC was the glutamine transporter SLC38A1, while the most downregulated gene was SLC1A7, the glutamate transporter, suggesting intramuscular glutamate-glutamine handling is significantly altered in sepsis.

**Figure 1.**
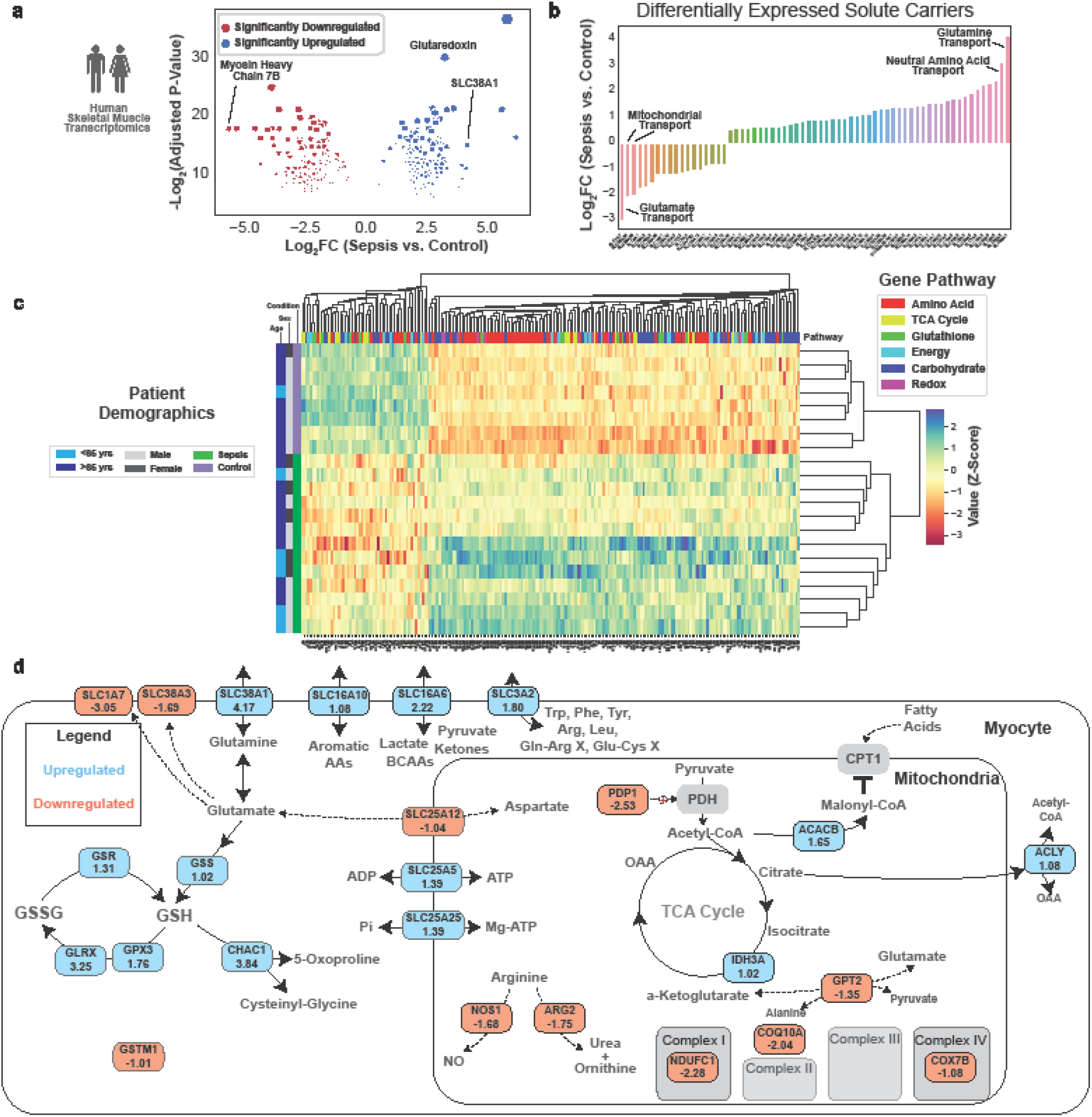
Human skeletal muscle transcriptomics reveals substantial metabolic reprogramming. (A) Volcano plot with significantly up and downregulated genes of septic and control patient skeletal muscle gene expression. (B) Barplot of differentially expressed solute carriers. (C) Clustermap of differentially expressed genes in one of six specified pathways. Patient identifiers are on the left most part of the figure, and gene classifiers are at the top portion of the figure. (D) Metabolic map of differentially expressed genes. All displayed genes have an adjusted p<0.05 from differential gene expression analysis.

Unsupervised clustering analysis with genes involved in the six pathways again demonstrated clear separation between septic and control patients (Fig 1C). There was no apparent sex-related clustering. Though 17% of patients (1/6) in the cluster more similar to the control muscle were younger than 65 years old, while 57% of patients (4/7) in the more opposite cluster were younger than 65, consistent with the finding that younger individuals have a more severe sepsis-induced muscle wasting phenotype^19^, there was no statically significant difference between age groups in this dataset.

Focusing on genes that are differentially expressed more than twofold (Fig 1D), we observed significant downregulation of electron transport genes, but upregulation of mitochondrial ATP transport. Glutamate transporters on the plasma membrane (SLC38A3 and SLC1A7) and mitochondrial GPT2, which mediates glutamate entry into the TCA cycle, were downregulated, while glutamine transporter SLC38A1, glutathione synthase (GSS), and multiple glutathione cycling genes were upregulated. In addition, transporters for branched-chain amino acids, aromatic amino acids, and arginine and cysteine were upregulated, providing a potential mechanism for degradation and release of amino acids in skeletal muscle. Together, our transcriptomics analysis highlighted a major reprogramming of amino acid handling, and impairment in electron transport chain function in skeletal muscle of sepsis patients.

### Moderate Cecal Ligation and Puncture Induces Body Weight Loss, Inflammation, and Reduction in Activity, Oxygen Consumption, and Carbon Dioxide Production

To further interrogate the metabolic mechanisms of sepsis, we performed a comprehensive characterization of our murine model of polymicrobial sepsis, cecal ligation & puncture (CLP)^20,21^. We determined that ligation of 0.5cm of cecum (Fig 2A) with a single puncture with a 23g needle, followed by sutures at two locations (Fig 2B-C) led to a one-week survival of approximately 68%, which approximates the lower end of survival rate estimates in humans (67-83%)^5,6^ (Fig 2D). On average this degree of CLP induced 20-25% reduction in body weight within 48 hours (Fig 2E). Septic mice also had widespread induction of numerous inflammatory cytokines, including key players IL-6, IL-10, and MCP-1 (Fig 2F). Metabolic cage studies demonstrated a significant reduction in voluntary activity after sepsis induction compared to mice that underwent a sham procedure (Fig 2G), coinciding with reduced water intake (Fig 2H), and reduced oxygen consumption (Fig 2I) and carbon dioxide production (Fig 2J). Based on these data, we performed endpoint metabolic studies at the 16-hour post-sepsis timepoint, where mice are reproducibly hypometabolic: voluntary activity was low and comparable between both sham and CLP mice, and systemic metabolism was substantially reduced in CLP mice.

**Figure 2.**
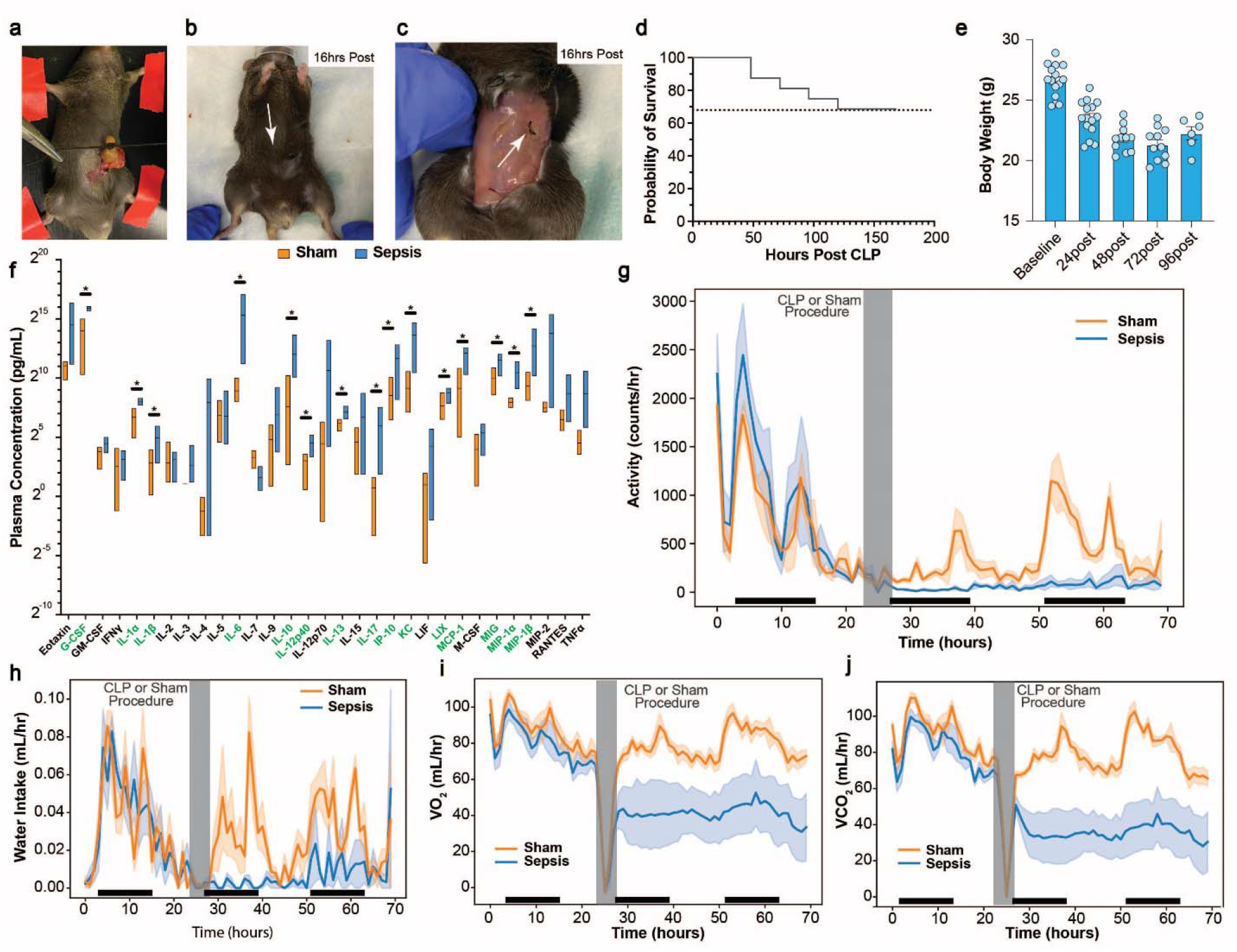
Murine model of sepsis, Cecal Ligation and Puncture, induces weight and activity loss, widespread inflammation, and significant reduction in energy expenditure. Photos of degree of ligation (A), superficial suture (B), and internal suture (C) during CLP. (D) Survival curve for degree of mortality. (E) Body weight during CLP challenge. (F) Plasma cytokines in septic and sham control animals. Metabolic cage data showing reduction in activity (G), water intake (H), VO_2_ (I), and VCO_2_ (J). Dark bars along the x-axis indicate the animal’s dark cycle. Asterisk in F indicates p-value from student’s t test < 0.05 between sepsis and sham groups.

### Plasma Metabolomics Reveals Systemic Metabolic Cycle Activity, Stress Hormone Activity, and Global Oxidative Stress

Untargeted and targeted liquid chromatography-mass spectrometry/mass spectrometry (LC-MS/MS)-based metabolomics were performed on plasma collected 16 hours post sepsis or sham surgery. Principal component analysis effectively distinguished the experimental groups with principal component 1 explaining 48.5% of the variance in the datasets (Fig 3A). Hippurate, which was administered as methenamine hippurate to prevent recurrent urinary tract infections^22,23^, was significantly downregulated in sepsis, while glutamine and classic inflammation-induced metabolite, itaconate^24^, were significantly upregulated (Fig 3B). Metabolites within the NAD biosynthetic pathway were significantly disturbed, with a notable 4-fold reduction in circulating nicotinamide and elevation of kynurenine (Fig 3C). Unsupervised clustering analysis revealed significant downregulation of carbohydrate metabolites, with upregulation of glutamine, stress-related metabolites including TCA, urea cycle, and xenobiotic metabolites (Fig 3D). The stress response during the tolerance phase of sepsis was apparent with 4-10 fold increases in cortisone, corticosterone, and itaconate (Fig 3E). Broadly, carnitines were also upregulated, providing evidence for a metabolic mechanism that supports ammonium clearance during increased systemic fatty acid oxidation (Fig 3F). Of the amino acids, glutamine elevation and glutamate and methionine suppression were the most notable signatures, consistent with the human transcriptomics data (Fig 3G). Consistent with a picture of systemic oxidative stress and mitochondrial dysfunction^25^, TCA intermediates were broadly reduced (Fig 3H). Further, the elevated benzoic acid and reduced hippurate is consistent with enhanced nitrogen clearance, possibly engaging this particular pathway due to gut microbial synthesis of these metabolites (Fig 3I).

**Figure 3.**
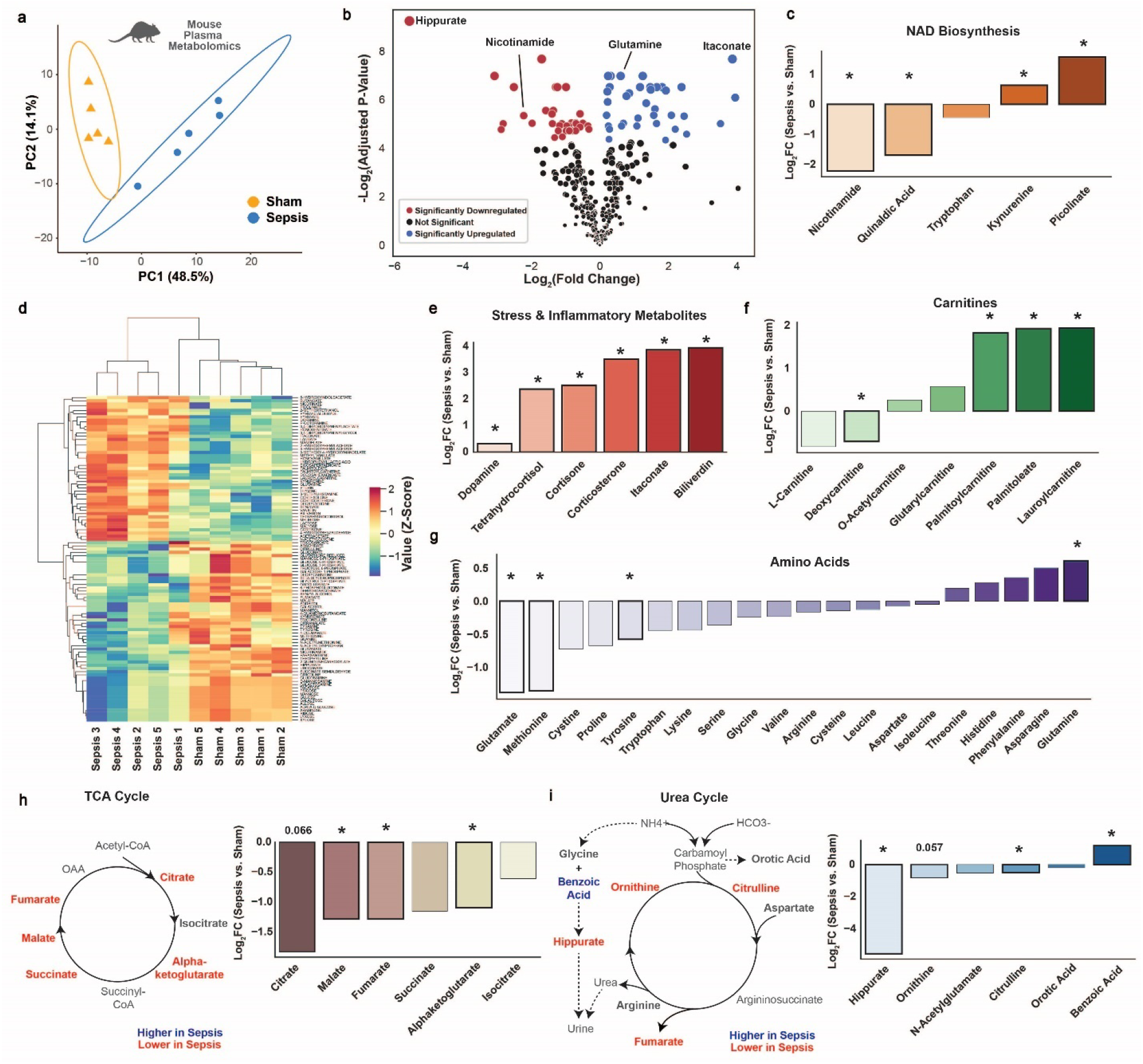
Plasma metabolomics demonstrates high degree of metabolic reprogramming at the systemic level. (A) Principal components analysis of plasma metabolomics. (B) Volcano plot, with differentially expressed metabolites (adjusted p<0.05) in red if downregulated, and blue if upregulated in septic mice. (C) Metabolites in the NAD Biosynthetic pathway. (D) Clustermap of all differentially expressed metabolites. Select differentially expressed metabolites related to inflammation/stress (E), carnitines (F), and amino acid metabolism (G). A metabolic map and associated bar plot of key metabolites in the tricarboxylic acid (TCA) cycle and related pathways (H), and in the urea cycle (I). An asterisk indicates a benjamani-hochberg adjusted p<0.05 for septic vs. sham animals.

### Tissue-Specific and Shared Metabolic States are Reprogrammed During Sepsis

To understand the unique and shared metabolic features of each organ system during pathogen-induced inflammation, we performed unsupervised hierarchical clustering of the metabolites in seven tissues of sham and septic animals (Fig 4A). We found that the same organs were clustered together while sham and septic groups were effectively distinguished, confirming that tissue-specific and sepsis-specific metabolic features are robust measurements for characterizing the disease in various organs. Interestingly, branched-chain amino acids (valine, leucine, and isoleucine) were highly enriched in the liver among other organs, and septic liver had even higher enrichment of these metabolites compared to the sham control. In addition, NADP+ level appeared highest in septic livers, which serves as a key signal for upregulation of NADPH synthetic pathways^26^. There appeared to be a muscle-specific set of metabolites including proline, lysine, alanine, and hexose-phosphate, consistent with both high free amino acid content of skeletal muscle and the lack of glucose-6-phosphatase in skeletal muscle, leading to high intracellular content of glucose-6-phosphate.

**Figure 4.**
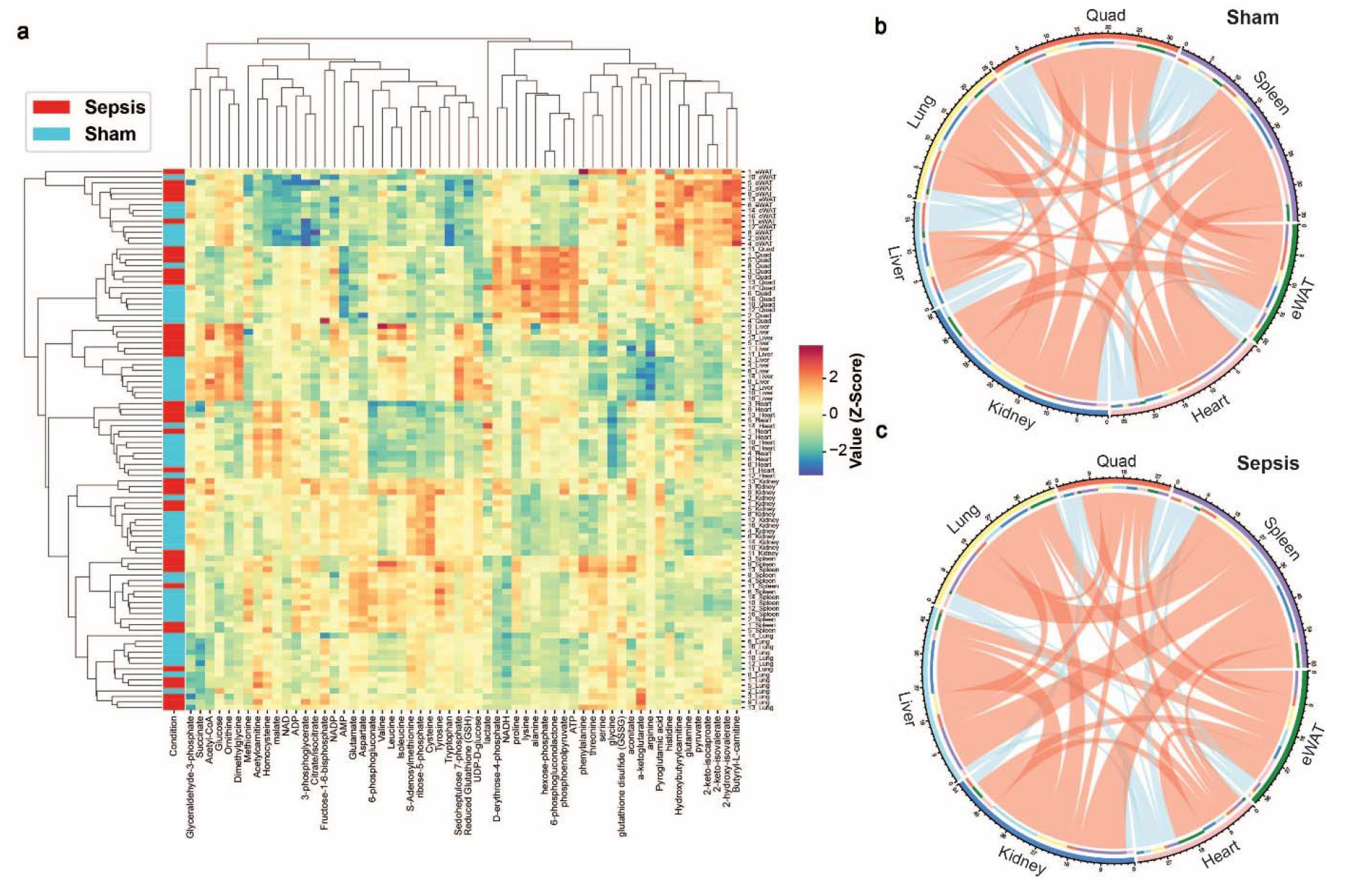
Shared and tissue-specific metabolic responses to sepsis. (A) Clustering heatmap shown for all measured intra-tissue metabolites in septic and sham control mice. Chord plot demonstrating significant correlations (p<0.05 for spearman rho correlation) of metabolites between seven tissues in sham (B) and septic (C) mice. Red lines indicate positive correlations, blue lines indicate negative correlations. The line thickness is dictated by the number of metabolites shared between tissues.

To understand the shared metabolic phenotype among different organs, we performed spearman rho correlations between pairs of all tissues (Fig 4B,C). In the sham condition, the liver had the fewest (18) significantly correlated metabolites, owing to its unique role in maintaining metabolic homeostasis. The metabolite with the most significant correlations in the control condition was S-adenosyl-methionine (Extended Data Fig 2A), providing evidence for its ubiquitous role in metabolic homeostasis through one-carbon metabolism at the whole-body level during rest. During sepsis, the relative number of correlations shared by cardiac and skeletal muscle dropped to a large degree, while the abdominal organs (spleen, kidney, liver) shared a much greater proportion of metabolites, in particular glutamate and glutamine (Extended Data Fig 2B). These data suggest unique metabolic states for heart and skeletal muscle, and a global coordination of key metabolic processes, particularly amino acid metabolism, in other organs in response to sepsis.

### Cellular Energy Status and Cytosolic and Mitochondrial Redox States Set Metabolic State in Liver and Other Organs

Next, we investigated whether tissue-specific redox balance and energy states were altered during sepsis. The AMP/ATP ratio, which is a barometer of the energy status of the tissue, was significantly upregulated in the liver, providing a driving force for the influx of metabolites for ATP provision in the liver (Fig 5A). In the liver and spleen, the 2GSH:GSSG ratio was significantly reduced in septic mice, suggesting these organs are particularly under redox stress during sepsis (Fig 5B). At the whole organ level, NAD/NADH ratio was increased only in the septic spleen (Extended Data Fig 3C). However, at the subcellular compartment level, NAD/NADH was altered in multiple organs by sepsis. For example, mitochondrial redox state, indicated by the glutamate:alpha-ketoglutarate ratio, was lower in the spleen (Extended Data Fig 3B), and cytosolic redox state, indicated by the lactate: pyruvate ratio, was lower in liver, spleen, and epididymal white adipose tissue (eWAT) (Fig 5C). The ADP/ATP ratio was increased modestly in the lung but tended to decrease in the spleen and heart, hinting at the existence of signals to reduce energy production in these two tissues (Extended Data Fig 3A).

**Figure 5.**
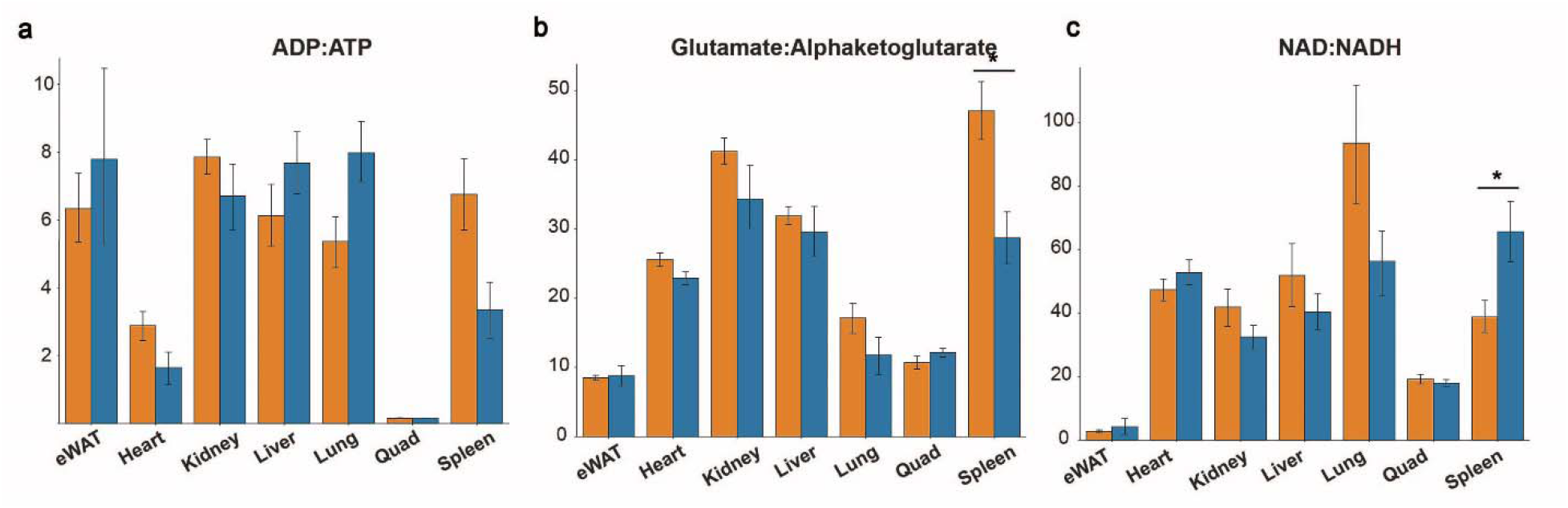
Sepsis induces global redox and energetic stress in tissues. (A) Tissue energy state ratios of ADP to ATP and AMP to ATP. (B) Redox ratios of reduced glutathione to glutathione disulfide (2GSH:GSSG) and (C), lactate to pyruvate. Asterisk indicates p<0.05 from a student’s t test between septic and sham animals.

### Glutamine Fuels Liver Glutathione Synthesis and TCA-Cycle Dependent Processes, While Skeletal Muscle Glutamine Metabolism is Shut Down

To understand how tolerance to sepsis reprograms systemic nutrient turnover and tissue-specific substrate utilization, we performed stable isotope infusions with ^2^H_7_-glucose and ^13^C_5_-glutamine. Both glucose and glutamine reached steady state enrichments by 110 minutes of the 120-minute infusion (Fig 6A,B). We observed that glucose turnover was significantly reduced (Fig 6C) while glutamine turnover was not altered in septic mice (Fig 6D). However, despite the lack of difference in whole-body glutamine turnover between sham-operated and CLP mice, we probed further to determine whether any differences in glucose metabolism may be present. As has been implied^27^, circulating glutamine contributed significantly more to *de novo* glutathione synthesis in the liver (Fig 6E), but was not upregulated in any other organs, demonstrating a muscle-liver glutamine-to-glutathione axis. Glutamine also served a more significant role in TCA anaplerosis in the kidney and the liver, as noted by the significant increases of the fractional contribution of glutamine to TCA cycle intermediates (Fig 6F). Meanwhile, glutamine contributed to the TCA cycle to a much lesser degree in the skeletal muscle and the spleen (Fig 6F).

**Figure 6.**
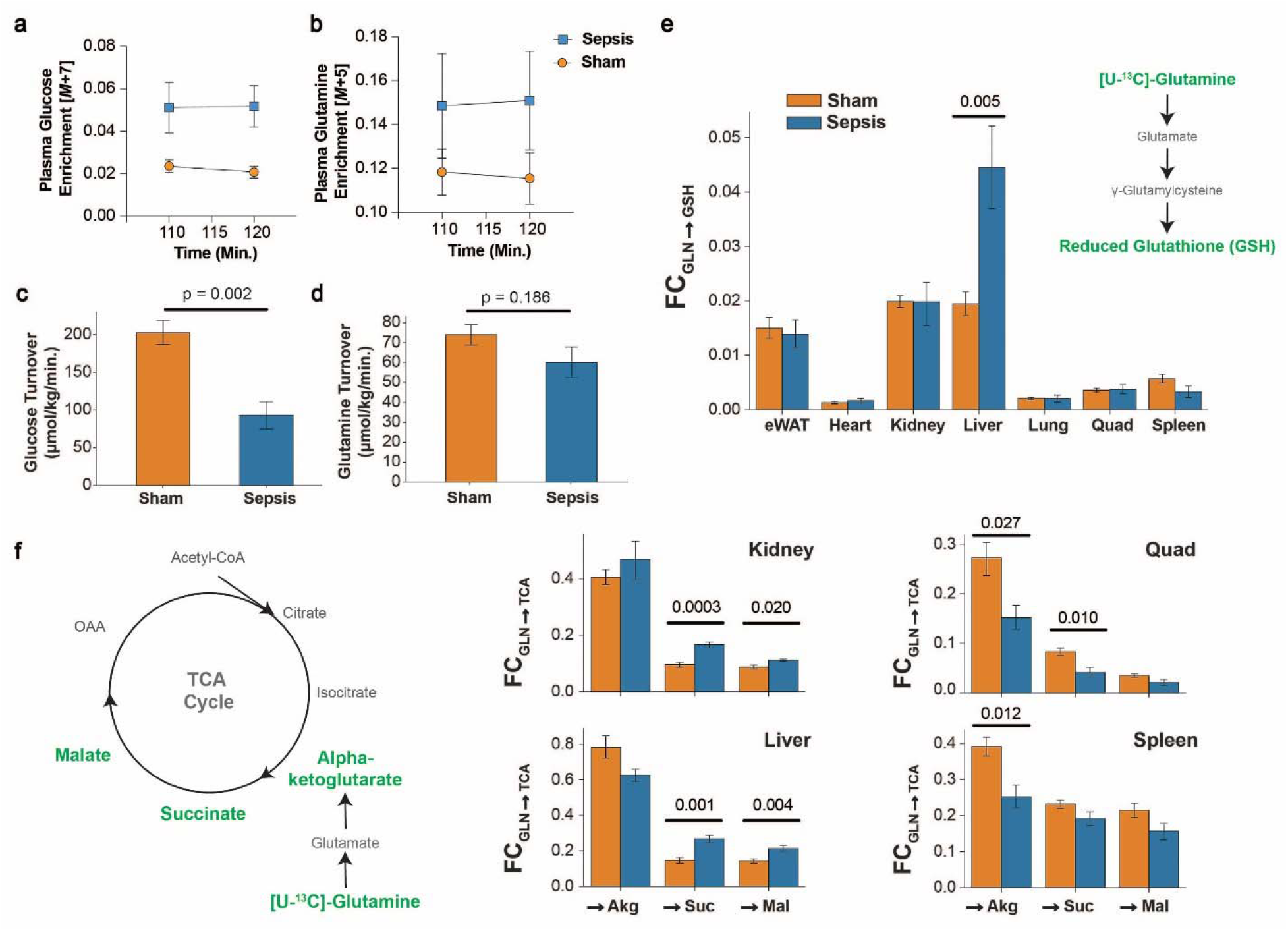
Stable isotope tracing with U-^13^C_5_-Glutamine reveals muscle-liver glutamine reprogramming. Fully labeled tracer enrichment for [1,2,3,4,5,6,6-^2^H_7_]-glucose (A), and [U-^13^C_5_]-glutamine (B) during the final ten minutes of the tracer infusion study. Whole body turnover of glucose (C) and glutamine (D). Fractional contribution of glutamine to glutathione (E), and to three intermediates of the TCA cycle (F). p<0.05 is deemed significant from a student’s t test between septic and sham animals.

## Discussion

The metabolic response to sepsis has long been studied and is still of great interest for guiding therapeutic management of critical illness and long-term morbidity^1^. However, many previously investigated pathways have not led to practice-changing clinical trials, and thus the field is at a stage where many of the well-studied metabolic pathways including glucose production^28–31^, lactate production^32,33^, and muscle proteolysis^34,35^ require greater insight and may require pivoting to the exploration of new metabolic pathways, to translate to standard-of-care precision medicine. The more recent advent of high-throughput biological omics techniques (transcriptomics, proteomics, metabolomics, and others) provides a timely platform to discover new metabolic mediators of the coordinated metabolomic response to sepsis. Numerous studies have utilized these techniques and provided unique insights into otherwise underappreciated pathways, most often on samples obtained solely from the plasma and/or from a single organ. However, few have combined these systems biology approaches with targeted, stable isotope tracer-based metabolic flux techniques, the gold-standard to elucidate changes in metabolite flux in plasma and in a comprehensive set of tissues. Thus, the goal of our study was to elucidate pathways of inter-organ metabolic crosstalk that define the metabolic response to sepsis, provide greater insight into previously characterized studies, and finally determine the metabolic mechanisms of the fate of glutamine at the systemic level.

A major difficulty with designing therapies to modulate systemic metabolism for critical illness is the timing of metabolite modulation. Steroids are among the few drugs given during acute sepsis that can improve survival. Consistent with this, endogenous steroids cortisone and hydroxycortisone were elevated in our untargeted metabolomic analysis, which contribute to enhanced skeletal muscle proteolysis. However, methenamine hippurate, a version of hippurate which also appeared in our analyses can effectively prevent recurrent urinary tract infections when administered prophylactically^22^ but is not effective at reducing illness when given acutely during sepsis. These two examples of applied metabolomics suggest that discovery techniques should provide suggestions, but targeted, mechanistic follow-up will be critical to determine how therapy can be designed to improve survival and reduce morbidity. Glutamine metabolism, which appeared in both our skeletal muscle transcriptomic and plasma metabolomic studies, is the example with which we followed up in a targeted manner.

Glutamine in the context of sepsis-induced critical illness has been investigated thoroughly^9,10,36–38^, and so far the few consensus opinions are that glutamine supplementation appears ineffective at reducing skeletal muscle proteolysis^39^, and that it should not be administered in critically ill patients^40^, especially in light of the evidence from the REDOXs randomized controlled trial, in which investigators demonstrated that glutamine supplementation worsened survival in patients with sepsis, though it should be kept in mind only nearly half of these patients had confirmed microbial identification, so the influence of viral versus bacterial versus fungal infections cannot be determined^2^. A key insight that has been largely missing from prior studies is an answer to: what is the fate of glutamine within the tissue/cell during critical illness? It is well-supported that glutamine is likely derived from skeletal muscle proteolysis^41^, but where does it go?

Our data supports a model by which skeletal muscle proteolysis fuels liver-mediated *de novo* glutathione synthesis, as well as liver amino acid reprogramming. These processes are dependent both on the redox and energy status of the organism, and in particular the status in the liver. When liver energy demand is high (indicated by an elevated AMP:ATP as in our study), substrate delivery can become limiting for energy and antioxidant-producing pathways. Thus, the most effective proteolyzers (young people as suggested by our data, and more muscular individuals as has been documented^19^) will support more energy production and antioxidant synthesis. Given data from the REDOXs trial that demonstrated that glutamine supplementation during critical illness increases mortality, however, increased glutamine supply appears detrimental. Thus, more effective strategies geared toward modulation of glutamine and glutathione pathways may reconsider targeting liver energy and redox states, more direct inhibition of proteolysis, or clearance of toxic metabolic byproducts of amino acid metabolism (nitrogen species). Our data support the idea that glutamine supplementation likely does not inhibit proteolysis by uptake and further metabolism, considering substantially lower glutamine incorporation into TCA cycle metabolites. Rather, it appears that amino acid metabolism is shut off in the muscle so that it can serve the sole purpose of releasing its critical amino acid stores.

A common conclusion from studies examining the metabolic response to organ damage is that there is systemic mitochondrial dysfunction, which in many cases may be based only on gene expression. By tracing glutamine’s incorporation into key TCA intermediates, we can understand how glutamine-supplied oxidation in the TCA cycle is altered mechanistically. We demonstrate that only in spleen and skeletal muscle does glutamine contribute significantly less to alpha-ketoglutarate, consistent with mitochondrial dysfunction or suppression in these tissues. In kidney and liver, however, the contribution of glutamine to the TCA intermediates succinate and malate is significantly higher in sepsis suggesting either a greater role for glutamine to TCA oxidation and/or impaired alternative (unlabeled) metabolite influx in the TCA cycle. In sum, the term mitochondrial dysfunction may not encompass the true complexity of tissue-specific mitochondrial metabolic reprogramming during sepsis.

Certain metabolic features of sepsis are shared among other various metabolic states, including starvation, ischemia-reperfusion injury (IRI), and several inborn errors of metabolism. In starvation, proteolysis ensues as glycogen stores are depleted, yet proteolysis rates appear to eventually slow^42^. Moreover, at least in human starvation, plasma glutamine is reduced while tyrosine and branched-chained amino acids are elevated, while in our experimental sepsis model glutamine was elevated, tyrosine was significantly reduced, and BCAAs tended to be reduced. The mechanisms for the shared and distinct regulation of amino acids during sepsis versus starvation remain to be elucidated. In IRI, elevated plasma succinate is a hallmark^43^, where we observed reduced succinate, possibly because the putative reduction in perfusion in our study may be less severe than experimental IRI. Our septic metabolome appears nearly opposite that of the mitochondrial disorder leading to mitochondrial encephalomyopathy lactic acidosis and stroke-like episodes (MELAS), which is dominated by reductive, rather than oxidative stress^25^. Though there are shared metabolic features of our experimental model of sepsis and numerous other systemic metabolic conditions, there appear metabolic mechanisms unique to each that need further investigation.

However, the metabolic response to sepsis appears strikingly similar to urea cycle disorders, in particular ornithine transcarbamylase deficiency^44^. Elevated benzoic acid in the setting of reduced urea cycle metabolites including citrulline and ornithine, as well as hippurate, which is a renally cleared nitrogen carrier illustrates an intriguing therapeutic target. Benzoic acid is FDA approved for the treatment of urea cycle disorders due to its ability to conjugate with the amino acid glycine and thus serve as a nitrogen carrier in the setting of excess ammonia/urea production. As increased plasma nitrogen content is associated with severity of sepsis,^45^ enhanced clearance may be an important therapeutic consideration. In addition, targeting nitrogen clearance rather than inhibition or supplementation of specific individual amino acids is plausibly higher yield, as key metabolic process to mount an immune response and maintain homeostasis can be enabled (but likely not enhanced). It remains to be determined whether impaired nitrogen clearance *per se* or something upstream is a main mediator of mortality.

Consistent with the observations that deadly immunotherapy-induced cytokine release syndrome appears in individuals with the most robust immune responses, we posit that supporting immune function, redox, and energy metabolism through glutamine may be detrimental, and support for these processes either by effective proteolysis or glutamine supplementation may support too robust of an inflammatory response. In addition, we provide an explanation for why glutamine supplementation likely will not improve skeletal muscle protein balance-glutamine is hardly incorporated into skeletal muscle metabolic pathways during sepsis. Rather, enhancing nitrogen clearance, which may be less likely to alter normal partitioning of key amino acid-regulated processes, represents an intriguing therapeutic possibility during sepsis. Our multi-omic analyses provide new insights into metabolic reprogramming during sepsis and suggest a better understanding of tissue-specific and cellular-compartment specific redox state, inter-tissue metabolic regulation, as well as the metabolic destiny of intratissue glutamine, which may hold promise as new metabolic modulation therapy during sepsis-induced critical illness.

## Acknowledgements

We gratefully acknowledge Dr. Gang Peng for conversations on statistical analyses, Ali Nasiri for assistance with metabolic cage studies, and the staff at the Yale Animal Resources Center (YARC) for their careful monitoring of the animals in this study. In addition, we thank Dr. Andrew Wang for guidance on performing cecal ligation and puncture. We thank the IOMIC Core at Yale for running the untargeted metabolomics. We are grateful to all members of the Perry lab and Rabinowitz lab for their helpful discussion.

B.P.L is supported under the National Institutes of Health Medical Scientist Training Program Training Grant T32GM007205. W.D.L is supported by NIH grant F32DK127843.

## Author Contributions

Conceptualization, B.P.L. and R.J.P.; Methodology, B.P.L., W.D.L., J.D.R., and R.J.P.; Investigation, B.P.L., W.D.L., W.Z., X.Z., R.C.G., and Z.L.; Writing – Original Draft, B.P.L.; Writing – Review & Editing, B.P.L, W.D.L, J.D.R., and R.J.P.; Funding Acquisition, R.J.P. and J.D.R.; Resources, R.J.P and J.D.R.; Supervision, R.J.P.

## Declaration of Interests

The authors declare no competing interests.

## Data and Code Availability

All data and code follow the FAIR principle as defined by the NIH Data Commons. Human RNA Transcriptomics can be found at GEO Accession Number GSE13205 at https://www.ncbi.nlm.nih.gov/geo/. Tissue and plasma metabolomic data with isotopic labeling can be found at. All Python and R code used to create the figures can be found on GitHub at https://github.com/BrooksLeitner/sepsismetabolism.

## Extended Data Figures

**Extended Data Figure 1.**
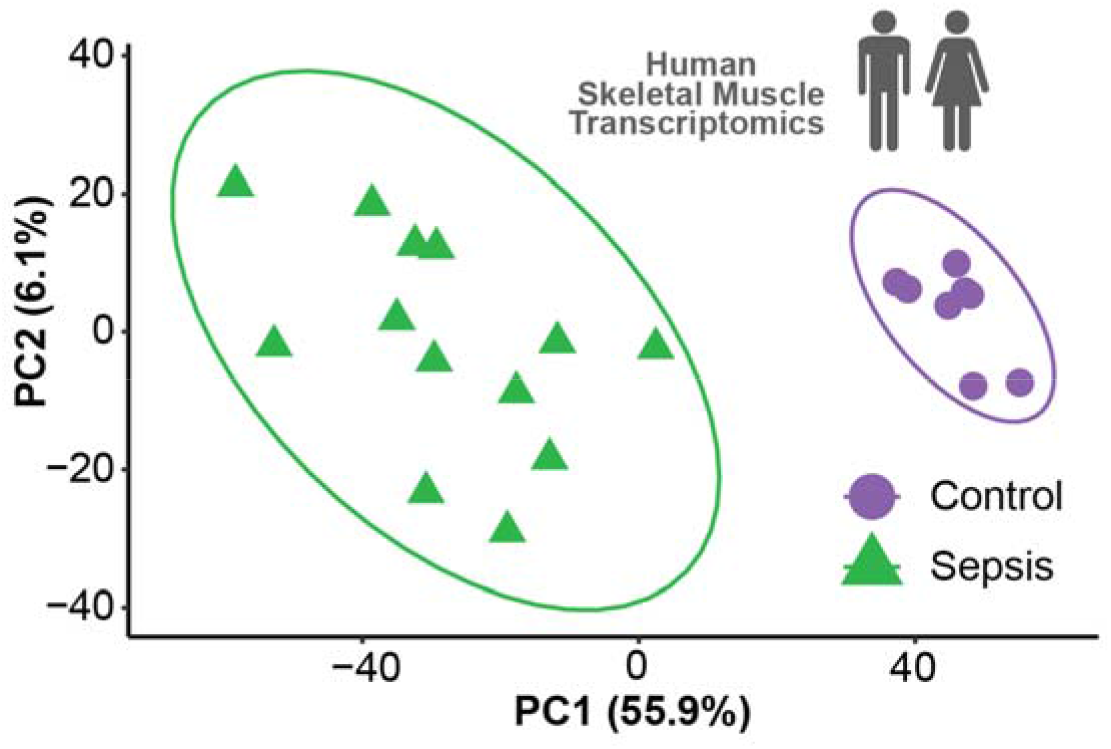
Principal components analysis of human skeletal muscle transcriptomics in patients with sepsis or control patients.

**Extended Data Figure 2.**
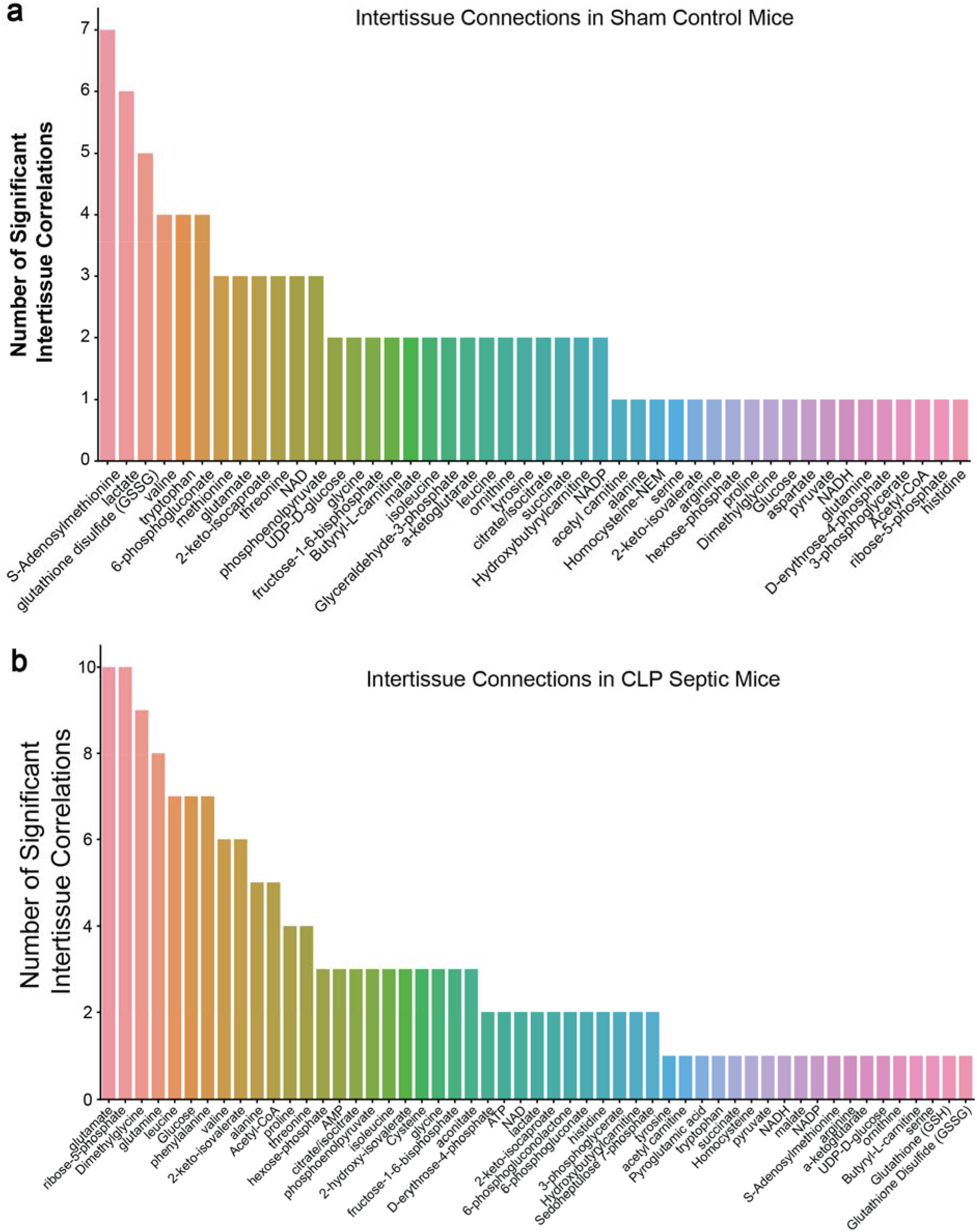
The cumulative number of significant correlations of intra-tissue metabolites between tissues in sham control mice (a), and septic mice that underwent cecal ligation and puncture (CLP).

**Extended Data Figure 3.**
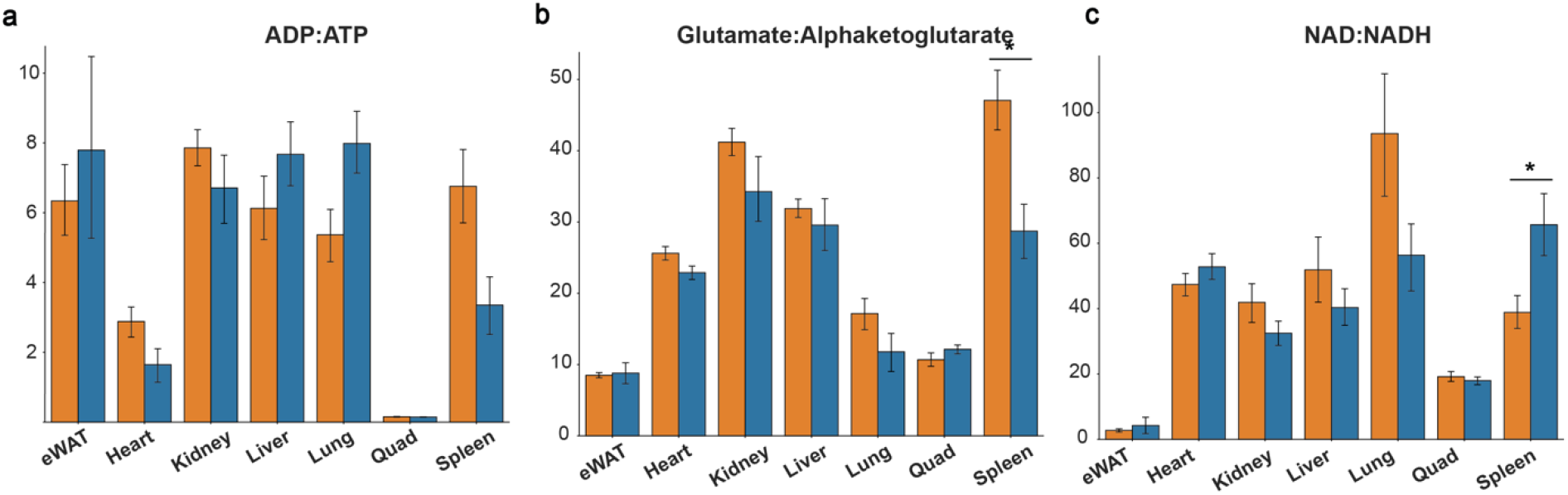
Intratissue redox and energetic ratios within tissues of mice with and without sepsis. Blue bars are septic mice and orange bars are sham control mice. *p < 0.05 by student’s t-test

**Extended Data Figure 4.**
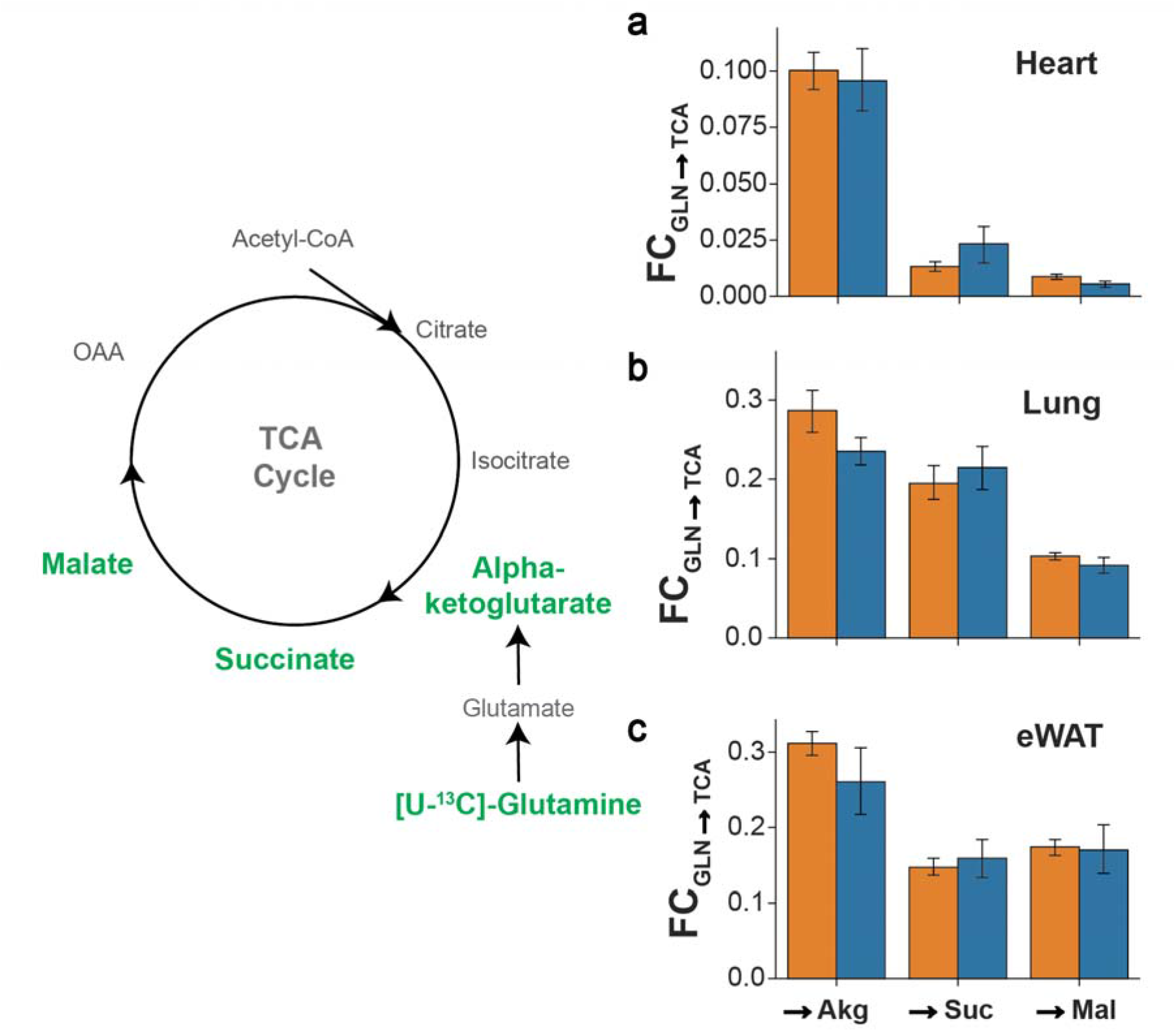
Fractional contribution of uniformly labeled ^13^C glutamine into tricarboxylic acid cycle (TCA) intermediates in the (a) heart, (b) lung, and (c) epidydimal white adipose tissue (eWAT). Akg = alphaketoglutarate, Suc = succinates, Mal = malate, GLN = glutamine. Blue bars are septic mice and orange bars are sham control mice.

## Materials and Methods

### Animals

All studies were approved by the Yale University Institutional Animal Care and Use Committee (protocols 2019-20290 and 2020-20149). 8-week-old male C57/Bl6J mice were maintained on standard rodent chow diet at room temperature in Yale Animal Research Center maintained housing. Cecal ligation and puncture (CLP) was performed under flow controlled (2.5%) isoflurane anesthesia. A 1 cm left of midline incision was made to both the cutaneous and peritoneal layers. The cecum was isolated and ligated at 0.5cm from the tip, then poked a single time with a 23g needle. Half an mL of PBS was injected into the peritoneal cavity before suturing closed the abdominal wall. Bupivacaine was applied between the abdominal wall and cutaneous skin layers, then a second suture closed the cutaneous portion of the abdomen. Sham control animals had the same procedure without the any manipulation of the cecum. Animals were randomly assigned to sepsis or sham control groups prior to studies. Food was removed from the cages and the mice were singly housed at the time of sham or CLP for survival, body weight, and untargeted plasma metabolomics studies. Mice were euthanized via isoflurane asphyxia followed by cervical dislocation 15-16 hours after CLP for the metabolomics and cytokine assays, and seven days after CLP for body weight and survival studies.

For stable isotope tracing, an intravenous catheter was placed through the jugular vein of the animal under isoflurane anesthesia, then tunneled subcutaneously with a thread at the top of the skin. Mice were singly housed after this surgery and were given carprofen in their drinking water for the following three days and checked daily to ensure recovery after surgery. Six days after catheterization surgery, mice were randomly allocated to CLP or sham control procedures, and food was removed from the cage at the time of surgery. Fourteen hours later, one end of the intravenous catheter was retrieved, flushed with 100 μL of normal saline, then attached to an infusion pump. U-^13^C_5_-Glutamine and ^2^H_7_-Glucose were co-infused for two hours (following a 3x prime for five minutes) continuous infusion. Tail blood was collected at minutes 110 and 120, and immediately centrifuged with supernatant plasma collected and stored in a −20°C freezer, to be used for plasma enrichment and steady state analyses. Then, mice were infused with 50 μL Euthasol (1:10 in PBS), blood from the inferior vena cava was immediately drawn and tissues collected and frozen in liquid nitrogen cooled Wollenberger tongs.

### Metabolic Cage Studies

Eight mice were singly housed three days prior to placement into the CLAMS apparatus from Columbus Instruments. After 24 hours in the metabolic cages, all mice were removed from the metabolic cages and either sham or CLP was performed within two hours. Mice were returned to the cage for 36 hours post-surgery with body weight determined at the end of the study.

### Cytokine Measurement

Blood from the inferior vena cava on the day of the stable isotope tracer studies was spun down for 30 seconds in a centrifuge and placed immediately in a −20C freezer. Samples were set to Eve Technologies (Details in the Key Resources Table) where the 31-Plex Cytokine Discovery Assay was run according to their instructions.

### Plasma Metabolomics

Ten mice were randomly subjected to CLP or Sham as described above. 20 μL of plasma was deproteinated and run on two liquid chromatography columns (Hypercarb and Reverse Phase) in both positive and negative modes for a total of 4 sample runs per study sample on LC-MS/MS. Targeted and untargeted metabolite peaks were curated using El-MAVEN software with the IROA library and KEGG database. ^2^H4-Taurine and ^2^H_8_-Phenylalanine were used as internal standards.

### Stable isotope tracing

Sixteen mice were randomly allocated to CLP or Sham as described above. Tracer was diluted in PBS to achieve infusion rates of 5.34 μmol/kg/min for [1,2,3,4,5,6,6-^2^H_7_]-glucose and 10 μmol/kg/min for [U-^13^C_5_]-glutamine ~2.5 hours prior to infusion. Mice were weighed immediately prior to infusion and a Harvard Syringe Pump was set to provide the correct infusion rate based on the animal’s body weight. Two mice in the septic group did not survive the infusion study, consistent with our survival percentage, and thus there were six mice in the CLP group and eight in the Sham procedure group.

At minutes 110 and 120 blood was drawn from the tail vein via tail massage and collected into capillary tubes, then immediately spun down in heparin-coated tubes, and the plasma was transferred to an Eppendorf for immediate placement into a −20C freezer. Mice were then euthanized and tissues were collected within 75 seconds in the following order after blood was drawn from the inferior vena cava: skeletal muscle, liver, kidney, spleen, epididymal white adipose tissue, heart, lung. Tissues once removed were immediately submerged in liquid nitrogen after clamping with pre-cooled Wollenberger clamps.

Tissues were ground to a fine powder under liquid nitrogen with a mortar and pestle. Ground tissue was then weighed (~ 20 mg) and mixed with −20°C 40:40:20 methanol:acetonitrile:water (extraction solvent) at a concentration of 25 mg/mL. Extract was then vortexed and centrifuged twice at 16,000 x g for 20 min at 4°C before the final supernatant was transferred to LC-MS tubes for analysis.

### Metabolite Measurement by LC-MS

The mass spectrometry analysis of polar metabolites was performed using Orbitrap Exploris 480 mass spectrometer (Thermo Fisher Scientific) coupled with hydrophilic interaction chromatography (HILIC). An XBridge BEH Amide column (150 mm × 2.1 mm, 2.5 μM particle size, Waters, Milford, MA) was used. The gradient was solvent A (95%:5% H2O:acetonitrile with 20 mM ammonium acetate, 20 mM ammonium hydroxide, pH 9.4) and solvent B (100% acetonitrile) 0 min, 90% B; 2 min, 90% B; 3 min, 75%; 7 min, 75% B; 8 min, 70% B, 9 min, 70% B; 10 min, 50% B; 12 min, 50% B; 13 min, 25% B; 14 min, 25% B; 16 min, 0% B, 20.5 min, 0% B; 21 min, 90% B; 25 min, 90% B. The flow rate was 150 μL/min with an injection volume of 5 μL and a column temperature of 25°C. The MS scans were in negative ion mode with a resolution of 480,000 at m/z 200. The automatic gain control (AGC) target was 1 × 10^6^ and the scan range was m/z 70-1000.

Data were analyzed using the El-MAVEN (Elucidata) software.^46^ Natural 13C and 2H abundances were corrected using AccuCor2 package in R.^47^

### Human Skeletal Muscle Transcriptomics

Data were accessed from GSE13205 using GEOquery package in R. Genes were filtered for low counts, then differential gene expression analysis was performed using Limma. Principal components analysis was performed in R, and then the differentially expressed gene list (all values with a benjamani-hochberg corrected p-value of less than 0.05) were exported to a CSV file. Gene set enrichment analysis was performed using EnrichR ^48^ (details in Key Resources Table), and HumanCyc 2016 pathways were selected due to their emphasis on metabolic pathways.

The volcano plot, bar plot, and clustering heatmap were created and generated in Python using the Seaborn package (details in Key Resources Table). The significantly differentially expressed genes were filtered using gene lists related to amino acid metabolism, the TCA cycle, glutathione metabolism, energy metabolism, carbohydrate metabolic and redox pathways. These gene lists were obtained from Reactome and MSigDB ^49,50^.

### Statistical Analysis

#### Human Skeletal Muscle Transcriptomics

GSE13205 was accessed using the R package GEOquery, and raw counts were filtered to remove genes that were not expressed in all samples, then log_2_(counts) transformed. A linear model was fit to the data using Limma, then fold change calculations in patients with sepsis versus control were calculated, with a Benjamani-Hochberg false discovery rate computed as the adjust p-value. In genes that were represented more once, the read with the higher logFC was selected.

A PCA plot was computed in R on filtered gene expression data and plotted using ggplot. The clustermap, barplot, and volcano plot were all created in Python using the Seaborn package on data that met the adjusted p-value <0.05, without a log2FC filter. For the metabolic gene expression map, differentially expressed genes identified in the clustermap were further filtered to those genes with a logFC >1.

#### Metabolic Cage Studies

A running filter down sampled the data to every hour and were plotted as means and SEMs using the Seaborn Python package.

#### Cytokine Measurement

Cytokines were quantified by Eve Technologies (Details in the Key Resources Table) 31-Plex Cytokine Discovery Assay, and a student’s t-test was used to compare sepsis vs. sham, with a p-value <0.05 deemed significant.

#### Plasma Metabolomics

2,531 unique metabolites were annotated. The fraction of each metabolite to the sum of all metabolites was computed for each metabolite, then log_2_(fraction of total) values were computed. Student’s t tests were performed on all metabolites between CLP and Sham, with a Benjamani-Hochberg p-value correction applied in Python (Code is Available, see details in the ***Data and Code Availability*** section). Differentially expressed metabolites were defined as an adjusted p-value <0.05, with no logFC threshold, as these analyses were used as a discovery analysis.

The PCA plot was generated in R. Pathway enrichment analysis was performed using MetaboAnalyst V5 ^51^ web tool. The volcano plot, bar plots, and clustering heatmap were created and generated in Python using the Seaborn package (details in Key Resources Table).

#### Plasma and Tissue Isotope Tracing

Metabolite peaks were background corrected using AccuCor2^47^, then converted to mass distribution vectors, such that each isotopologue was set as a fraction of the sum of the total isotopologues for each molecule. Pool size measurements, which reflect total molecular peak abundance, were used for calculating redox and energy status ratios. Student’s t tests were performed for CLP vs. Sham in GraphPad Prism, with a p-value <0.05 considered significant.

Turnover was calculated using the tracer to sum ratio method^52^. Binomial expansions were used to obtain natural abundance and tracer enrichment (98% for glucose, 99% for glutamine) mass distribution vectors (MDVs) of deuterium (0.0156%) and of ^13^C (1.1%) in glutamine^53,54^.

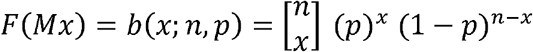

Where F is the frequency of an isotopologues, n is the number of potentially labeled atoms in the molecule, x is the number of actually labeled atoms in the molecule, and p is the probability of there being label in the atom (which corresponds either to the tracer enrichment or natural abundance enrichment). We devised a Python function that takes the MDVs of natural abundance (background), tracer, and mixed sample (plasma) enrichments, and computes a linear regression that resolves the estimate contribution from all three inputs to calculate turnover, which is available at the GitHub webpage listed in the ***Data and Code Availability*** section.

As we infused with a highly deuterated compound, which may be subject to primary and secondary isotope effects^55–58^, only ^13^C-labeled isotopologues were used for each metabolite in tissues. This method also minimizes the confounding influence of secondary tracers due to deuterium incorporation after metabolite flux through various pathways.

Fractional contribution (FC) analyses were performed using uniformly ^13^C-labeled plasma glutamine enrichment as the assumed only ^13^C labelled precursor. Atom percent excess (APE) was calculated as described^59,60^ from MDVs of plasma glutamine, and each specified tissue metabolite.

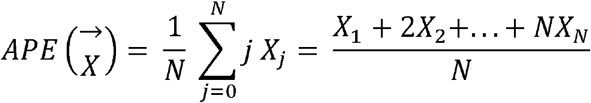

Where 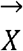 is the MDV of a molecule, N is the number of potentially labelled atoms in the molecule, and X is the isotopologue. This calculation of APE takes into consideration all labelled isotopologues of each metabolite and provides a weight that increases as the isotopologues is more highly labeled, and then is taken relative to the number of carbons, and also accounts for *in vivo* label dilution. For example, using the precursor-product relationship, FC_Gln→GSH_ described the calculation of (Tissue Glutathione APE) / (Plasma Glutamine APE). We devised a Python program that calculates APE for any metabolite given its MDV, which is available at the GitHub webpage listed in the Code Availability section.

## Data and Code Availability

Human skeletal muscle data are publicly available at GEO (GSE13205), mouse plasma metabolomics data and plasma & tissue fluxomic data will be available online on the date of publication. All original code has been deposited at GitHub (https://github.com/BrooksLeitner/sepsismetabolism). Any additional information required to reanalyze the data reported in this paper is available from the corresponding author upon request.

## Notes

### Competing Interest Statement

The authors have declared no competing interest.

## References

1. Deutschman, C. S. et al. The Surviving Sepsis Campaign: Basic/Translational Science Research Priorities*. Critical Care Medicine 48, 1217–1232 (2020).

2. Heyland, D. et al. A Randomized Trial of Glutamine and Antioxidants in Critically Ill Patients. N Engl J Med 368, 1489–1497 (2013).

3. Sevransky, J. E. et al. Effect of Vitamin C, Thiamine, and Hydrocortisone on Ventilator-and Vasopressor-Free Days in Patients With Sepsis: The VICTAS Randomized Clinical Trial. JAMA 325, 742–750 (2021).

4. Wernerman, J. et al. Metabolic support in the critically ill: a consensus of 19. Critical Care 23, 318 (2019).

5. Evans, L. et al. Surviving sepsis campaign: international guidelines for management of sepsis and septic shock 2021. Intensive Care Med (2021) doi:10.1007/s00134-021-06506-y.

6. Mostel, Z. et al. Post-sepsis syndrome – an evolving entity that afflicts survivors of sepsis. Mol Med 26, (2019).

7. Slikke, E. C. van der, An, A. Y., Hancock, R. E. W. & Bouma, H. R. Exploring the pathophysiology of post-sepsis syndrome to identify therapeutic opportunities. EBioMedicine 61, (2020).

8. Belsky, J. B., Wira, C. R., Jacob, V., Sather, J. E. & Lee, P. J. A review of micronutrients in sepsis: the role of thiamine, l-carnitine, vitamin C, selenium and vitamin D. Nutrition Research Reviews 31, 281–290 (2018).

9. Gore, D. & Wolfe, R. Metabolic response of muscle to alanine, glutamine, and valine supplementation during severe illness. Journal of Parenteral and Enteral Nutrition 27, 307–314 (2003).

10. Heyland, D. K. et al. Glutamine and Antioxidants in the Critically Ill Patient. Journal of Parenteral and Enteral Nutrition 39, 401–409 (2015).

11. Hasselgren, P.-O., Talamini, M., James, J. H. & Fischer, J. E. Protein Metabolism in Different Types of Skeletal Muscle During Early and Late Sepsis in Rats. Archives of Surgery 121, 918–923 (1986).

12. McGuinness, O. P. THE IMPACT OF INFECTION ON GLUCONEOGENESIS IN THE CONSCIOUS DOG. Shock 2, 336–343 (1994).

13. Meinz, H., Lacy, D. B., Ejiofor, J. & McGuinness, O. P. ALTERATIONS IN HEPATIC GLUCONEOGENIC AMINO ACID UPTAKE AND GLUCONEOGENESIS IN THE ENDOTOXIN TREATED CONSCIOUS DOG. Shock 9, 296–303 (1998).

14. Rhee, C. et al. Prevalence, Underlying Causes, and Preventability of Sepsis-Associated Mortality in US Acute Care Hospitals. JAMA Network Open 2, e187571–e187571 (2019).

15. Ganeshan, K. et al. Energetic Trade-Offs and Hypometabolic States Promote Disease Tolerance. Cell 177, 399–413.e12 (2019).

16. Gómez, H., Kellum, J. A. & Ronco, C. Metabolic reprogramming and tolerance during sepsis-induced AKI. Nat Rev Nephrol 13, 143–151 (2017).

17. Weis, S. et al. Metabolic Adaptation Establishes Disease Tolerance to Sepsis. Cell 169, 1263–1275.e14 (2017).

18. Fredriksson, K. et al. Dysregulation of Mitochondrial Dynamics and the Muscle Transcriptome in ICU Patients Suffering from Sepsis Induced Multiple Organ Failure. PLOS ONE 3, e3686 (2008).

19. Hart, D. W. et al. Determinants of Skeletal Muscle Catabolism After Severe Burn. Annals of Surgery 232, 455–465 (2000).

20. Dejager, L., Pinheiro, I., Dejonckheere, E. & Libert, C. Cecal ligation and puncture: the gold standard model for polymicrobial sepsis? Trends in Microbiology 19, 198–208 (2011).

21. Rittirsch, D., Huber-Lang, M. S., Flierl, M. A. & Ward, P. A. Immunodesign of experimental sepsis by cecal ligation and puncture. Nat Protoc 4, 31–36 (2009).

22. Lee, B. S. B., Bhuta, T., Simpson, J. M. & Craig, J. C. Methenamine hippurate for preventing urinary tract infections. Cochrane Database of Systematic Reviews (2012) doi:10.1002/14651858.CD003265.pub3.

23. Lees, H. J., Swann, J. R., Wilson, I. D., Nicholson, J. K. & Holmes, E. Hippurate: The Natural History of a Mammalian–Microbial Cometabolite. J. Proteome Res. 12, 1527–1546 (2013).

24. Chen, L.-L. et al. Itaconate inhibits TET DNA dioxygenases to dampen inflammatory responses. Nat Cell Biol 1–11 (2022) doi:10.1038/s41556-022-00853-8.

25. Sharma, R. et al. Circulating markers of NADH-reductive stress correlate with mitochondrial disease severity. J Clin Invest 131, (2021).

26. Chen, L. et al. NADPH production by the oxidative pentose-phosphate pathway supports folate metabolism. Nat Metab 1, 404–415 (2019).

27. Cruzat, V. F. et al. Oral free and dipeptide forms of glutamine supplementation attenuate oxidative stress and inflammation induced by endotoxemia. Nutrition 30, 602–611 (2014).

28. Jahoor, F. et al. Role of insulin and glucose oxidation in mediating the protein catabolism of burns and sepsis. American Journal of Physiology-Endocrinology and Metabolism 257, E323–E331 (1989).

29. Maitra, S. R., Wojnar, M. M. & Lang, C. H. ALTERATIONS IN TISSUE GLUCOSE UPTAKE DURING THE HYPERGLYCEMIC AND HYPOGLYCEMIC PHASES OF SEPSIS. Shock 13, 379–385 (2000).

30. Vary, T. C. et al. Pharmacological reversal of abnormal glucose regulation, BCAA utilization, and muscle catabolism in sepsis by dichloroacetate. J Trauma 28, 1301–1311 (1988).

31. Vary, T. C., Drnevich, D., Jurasinski, C. & Brennan, W. A. Mechanisms regulating skeletal muscle glucose metabolism in sepsis. Shock 3, 403–410 (1995).

32. Cakir, E. & Turan, I. O. Lactate/albumin ratio is more effective than lactate or albumin alone in predicting clinical outcomes in intensive care patients with sepsis. Scand J Clin Lab Invest 81, 225–229 (2021).

33. Garcia-Alvarez, M., Marik, P. & Bellomo, R. Sepsis-associated hyperlactatemia. Critical Care 18, 503 (2014).

34. Gore, D. C. & Wolfe, R. R. Hemodynamic and metabolic effects of selective β1 adrenergic blockade during sepsis. Surgery 139, 686–694 (2006).

35. Tiao, G. et al. Energy-ubiquitin-dependent muscle proteolysis during sepsis in rats is regulated by glucocorticoids. J. Clin. Invest. 97, 339–348 (1996).

36. Gianotti, L., Alexander, J. W., Gennari, R., Pyles, T. & Babcock, G. F. Oral Glutamine Decreases Bacterial Translocation and Improves Survival in Experimental Gut-Origin Sepsis. Journal of Parenteral and Enteral Nutrition 19, 69–74 (1995).

37. Noguchi, Y., Howard James, J., Fischer, J. E. & Hasselgren, P.-O. Increased glutamine consumption in small intestine epithelial cells during sepsis in rats. The American Journal of Surgery 173, 199–205 (1997).

38. Smedberg, M. et al. Plasma glutamine status at intensive care unit admission: an independent risk factor for mortality in critical illness. Critical Care 25, 240 (2021).

39. Gore, D. & Wolfe, R. Glutamine supplementation fails to affect muscle protein kinetics in critically ill patients. Journal of Parenteral and Enteral Nutrition 26, 342–349 (2002).

40. van Zanten, A. R. H. & Elke, G. Parenteral glutamine should not be routinely used in adult critically ill patients. Clinical Nutrition 36, 1184–1185 (2017).

41. Cruzat, V., Macedo Rogero, M., Noel Keane, K., Curi, R. & Newsholme, P. Glutamine: Metabolism and Immune Function, Supplementation and Clinical Translation. Nutrients 10, 1564 (2018).

42. Perry, R. J. et al. Leptin Mediates a Glucose-Fatty Acid Cycle to Maintain Glucose Homeostasis in Starvation. Cell 172, 234–248.e17 (2018).

43. Chouchani, E. T. et al. Ischaemic accumulation of succinate controls reperfusion injury through mitochondrial ROS. Nature 515, 431–435 (2014).

44. Burrage, L. C. et al. Untargeted Metabolomic Profiling Reveals Multiple Pathway Perturbations and New Clinical Biomarkers in Urea Cycle Disorders. Genet Med 21, 1977–1986 (2019).

45. Pittiruti, M. et al. Determinants of Urea Nitrogen Production in Sepsis: Muscle Catabolism, Total Parenteral Nutrition, and Hepatic Clearance of Amino Acids. Archives of Surgery 124, 362–372 (1989).

46. Agrawal, S. et al. El-MAVEN: A Fast, Robust, and User-Friendly Mass Spectrometry Data Processing Engine for Metabolomics. in High-Throughput Metabolomics: Methods and Protocols (ed. D’Alessandro, A.) 301–321 (Springer, 2019). doi:10.1007/978-1-4939-9236-2_19.

47. Wang, Y., Parsons, L. R. & Su, X. AccuCor2: isotope natural abundance correction for dual-isotope tracer experiments. Lab Invest 101, 1403–1410 (2021).

48. Kuleshov, M. V. et al. Enrichr: a comprehensive gene set enrichment analysis web server 2016 update. Nucleic Acids Res 44, W90–W97 (2016).

49. Liberzon, A. et al. The Molecular Signatures Database Hallmark Gene Set Collection. cels 1, 417–425 (2015).

50. Subramanian, A. et al. Gene set enrichment analysis: A knowledge-based approach for interpreting genome-wide expression profiles. Proceedings of the National Academy of Sciences 102, 15545–15550 (2005).

51. Pang, Z. et al. MetaboAnalyst 5.0: narrowing the gap between raw spectra and functional insights. Nucleic Acids Research 49, W388–W396 (2021).

52. Kelleher, J. K. & Masterson, T. M. Model equations for condensation biosynthesis using stable isotopes and radioisotopes. American Journal of Physiology-Endocrinology and Metabolism 262, E118–E125 (1992).

53. Hellerstein, M. K. Relationship between precursor enrichment and ratio of excess M2/excess M1 isotopomer frequencies in a secreted polymer. Journal of Biological Chemistry 266, 10920–10924 (1991).

54. Hellerstein, M. K. & Neese, R. A. Mass isotopomer distribution analysis: a technique for measuring biosynthesis and turnover of polymers. American Journal of Physiology-Endocrinology and Metabolism 263, E988–1001 (1992).

55. de Graaf, R. A., Thomas, M. A., Behar, K. L. & De Feyter, H. M. Characterization of Kinetic Isotope Effects and Label Loss in Deuterium-Based Isotopic Labeling Studies. ACS Chem. Neurosci. 12, 234–243 (2021).

56. Millard, P., Portais, J.-C. & Mendes, P. Impact of kinetic isotope effects in isotopic studies of metabolic systems. BMC Systems Biology 9, 64 (2015).

57. Westheimer, F. H. The Magnitude of the Primary Kinetic Isotope Effect for Compounds of Hydrogen and Deuterium. Chem. Rev. 61, 265–273 (1961).

58. Dale, H. J. A., Leach, A. G. & Lloyd-Jones, G. C. Heavy-Atom Kinetic Isotope Effects: Primary Interest or Zero Point? J. Am. Chem. Soc. 143, 21079–21099 (2021).

59. Buescher, J. M. et al. A roadmap for interpreting 13C metabolite labeling patterns from cells. Current Opinion in Biotechnology 34, 189–201 (2015).

60. Fernández-García, J., Altea-Manzano, P., Pranzini, E. & Fendt, S.-M. Stable Isotopes for Tracing Mammalian-Cell Metabolism In Vivo. Trends in Biochemical Sciences 45, 185–201 (2020).

